# Soft Tissue Mechanics in Hip Distraction after Total Knee Arthroplasty: A Finite Element Analysis

**DOI:** 10.1101/2024.02.13.580129

**Authors:** Sophia Soehnlen, Sara Sadeqi, Yogesh Kumaran, Sudharshan Tripathi, Ryan K. Jones, David H. Sohn, Vijay K. Goel

## Abstract

**INTRODUCTION:** Improvement in diagnostic and surgical techniques in hip arthroscopy have led to a surge in hip distraction procedures over the recent years with the predicted annual frequency being four out of every 10,000 orthopedic procedures in 2017. Due to the large traction force required to achieve the appropriate joint spacing intra-operatively, an emergence of traction-related neurological and soft tissue injuries have surfaced. Pre-existing hip joint pathologies and surgical procedures disrupt the biomechanical stability of the joint and significantly increase the risk of iatrogenic damage. Furthermore, patients with total knee arthroplasties are often subject to intra-articular ligament releases, leading to reduced stability; however, it is not well understood how this may impact their outcomes of hip arthroscopic procedures. The current study aims to investigate the biomechanical behavior of various instrumented knee joints subjected to traction forces to aid clinical understanding and advancements of hip arthroscopy techniques.

**METHODS:** A validated finite element (FE) model of the pelvis and lower extremity was developed from computed tomography (CT) scans of a healthy 45-year-old female. Three different models were assembled according to different TKA techniques performed: Bi-Cruciate Retaining (BCR) model, Posterior-Cruciate Retaining (PCR) model, and Posterior Stabilized (PS) model. The BCR model is noted by retaining all native ligaments of the knee joint (ACL, PCL, MCL, and LCL), whereas the PCR model was subject to ACL removal and the PS model required ACL and PCL removal (Figure 1). The pelvis was encastered to prevent translation under the traction forces as motion of the patient’s trunk is restrained, intraoperatively. To simulate the loading condition of hip distraction, an axial force was coupled to the distal fibula and tibia and incrementally increased from 100N to 500N. Joint spacing and ligament strain in the hip and knee joint were analyzed to assess the effects of traction forces.

Figure 1.
Meshed parts of the A.) Posterior Cruciate retaining total knee replacement system and the B.) Bi-cruciate retaining total knee replacement system.

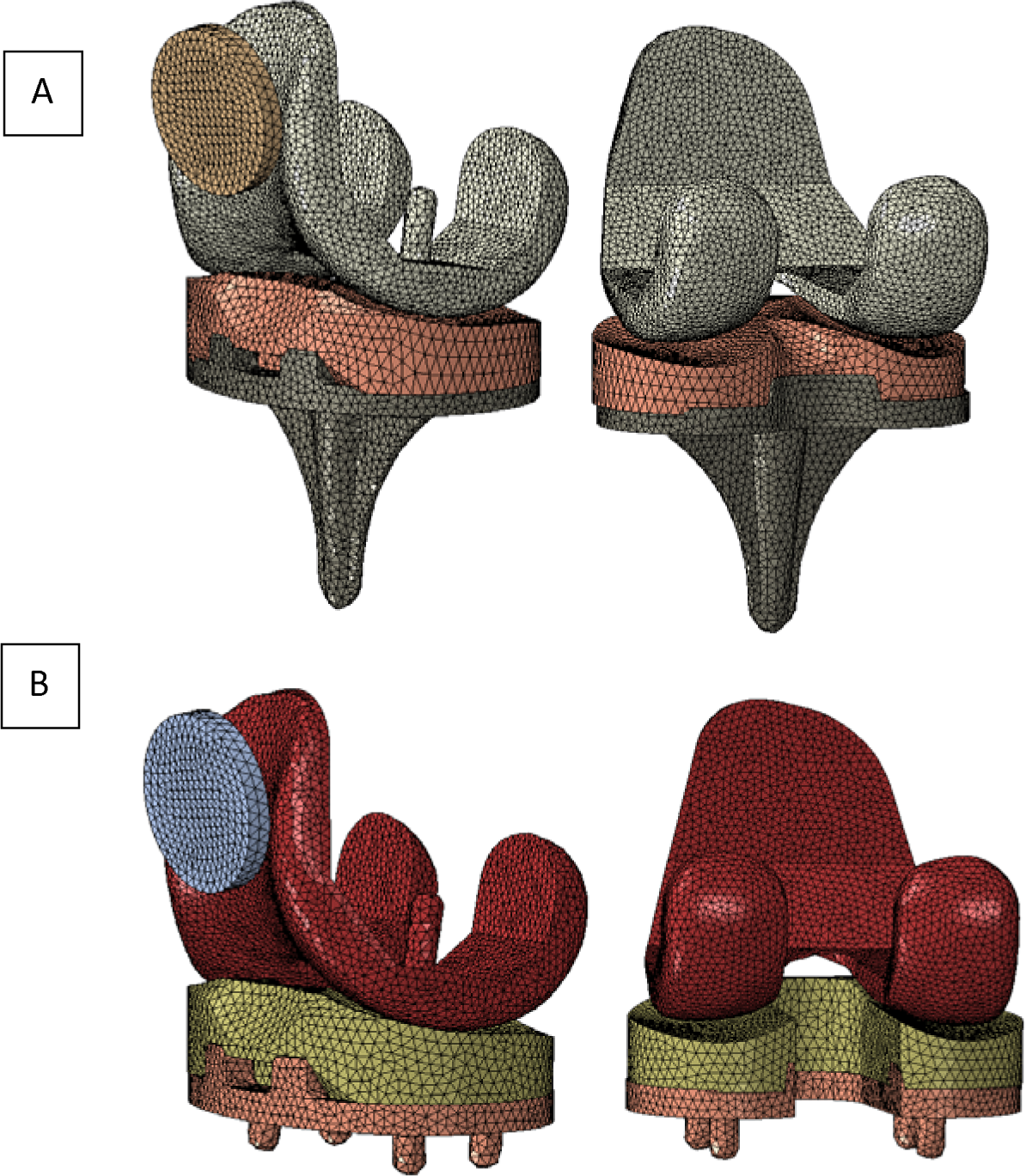

**RESULTS:** The medial and lateral compartment stiffness of the knee joint was analyzed under hip distraction for the three different TKA scenarios. The BCR model displayed the greatest average knee complex stiffness. Release of the ACL resulted in a larger decrease of stiffness compared to release of the PCL. There was no change in forces required for hip distraction as result of changes in the knee joint stiffness (Figure 3). The PCR and PS models were subject to excess knee joint distraction that exceeded 12 mm and ligament strain greater than 20% before adequate hip joint distraction of 10 mm was achieved. The BCR model remained below 10 mm of knee distraction and 15% ligament strain at 10 mm of hip joint distraction.

**DISCUSSION:** Our study reveals patients undergoing hip distraction with a prior TKA may experience increased soft tissue damage or iatrogenic dislocation due to reduced knee joint stability. The PCR and PS models outline a trend suggesting patients who have undergone ligament sacrificing TKAs experience large reductions in knee joint stability, causing strain levels that are indicative of soft tissue injury. The BCR TKA was indicated to be the safest under the distraction conditions as joint spacing and strain levels were largely reduced comparatively; however, when surpassing 10 mm of knee joint distraction at forces greater than 350 N, the strain levels in the ACL suggest minor injury may occur.

## Introduction

Hip distraction is a versatile surgical technique employed in minimally invasive hip arthroscopies for treating intra- and extraarticular pathologies and in a wide range of other interventions including joint preservation surgeries, management of hip dysplasia, treatment of avascular necrosis, fracture reduction and fixation, external fixation for pelvic injuries, and conservative management of hip conditions [add cite]. Recent advancements in hip arthroscopy, including novel instrumentation and refined surgical techniques, have not only increased the frequency of these procedures to four per 10,000 orthopedic interventions in 2017 [1], but also significantly reduced surgical complications and expedited patient recovery, presenting a marked improvement over traditional open surgical methods [2, 3]. During the procedure, achieving adequate joint visualization necessitates mild dislocation, achieved through the application of traction. Distraction devices employ a consistent biomechanical method, wherein the patient’s foot is secured in a boot connected to a gliding mechanism. This setup facilitates the application of traction forces, pulling the boot—and consequently the leg—away from the patient, generating an axial force through the lower extremity and joints [4, 5].

Due to the large traction force provided to achieve the appropriate joint spacing, an emergence of distraction-related injuries has surfaced that lead to neurologic and soft tissue damage [3, 6, 7]. Factors such as body weight, sex differences, and bone morphology play a crucial role in determining the necessary traction force for ideal joint spacing [8–10]. Moreover, pre-existing conditions and previous surgeries of the knee and hip joints can compromise the biomechanical stability of the joint, elevating the risk of iatrogenic damage during hip distraction [11]. A particular concern arises in patients with prior total knee arthroplasty (TKA), where the removal of the cruciate ligaments diminishes knee joint stability post-operatively [12, 13]. In response, the bicruciate sparing TKA technique has been introduced to preserve the biomechanical characteristics of a healthy knee, enhancing post-operative outcomes [14]. Nevertheless, the biomechanical integrity of the knee joint complex may still be at risk during hip distraction, regardless of the surgical technique employed.

To date, the mechanical behavior of the knee joint subjected to varying traction forces during hip arthroscopy in patients with a history of TKA remains unexplored. This gap in knowledge represents a critical barrier to optimizing surgical outcomes and minimizing the risk of iatrogenic injury. Our study seeks to bridge this gap by quantifying the displacement and strain exerted on the knee joint and its ligaments during hip distraction, specifically after different TKA techniques. Through a comprehensive parametrized finite element (FE) analysis, we aim to provide surgeons with detailed insights into the biomechanics of hip distraction, thereby informing safer surgical practices and enhancing patient care.

## Methods

### Model Development

A previously validated finite element (FE) model of the lower extremity was utilized in this study [15]. The bony and main ligament geometries of a 45-year-old female were obtained from computed tomography and magnetic resonance imaging, respectively. Mimics v15 (Materialise, Leuven, Belgium) was used for image reconstruction and imported into Geomagic Studio (3DSystems, Morrisville, North Carolina, USA) to smooth the geometries. Three-dimensional geometries for TKA instrumentation were created using Solidworks computer aided design (Dassault Systèmes, Vélizy-Villacoublay, France). Blender (Blender foundation) was utilized to perform Boolean functions to remove the bony features of the 3D geometry where the implant was placed. Following a previous study [15], the TKA was implanted under guidance of an orthopedic surgeon.

The hip capsule (Figure 2) was created in SolidWorks to fit the geometry of the model based on capsule attachment points noted by Stewart et al [16]. The capsule thickness fell within the ranges listed in literature [17].

**Figure 2.**
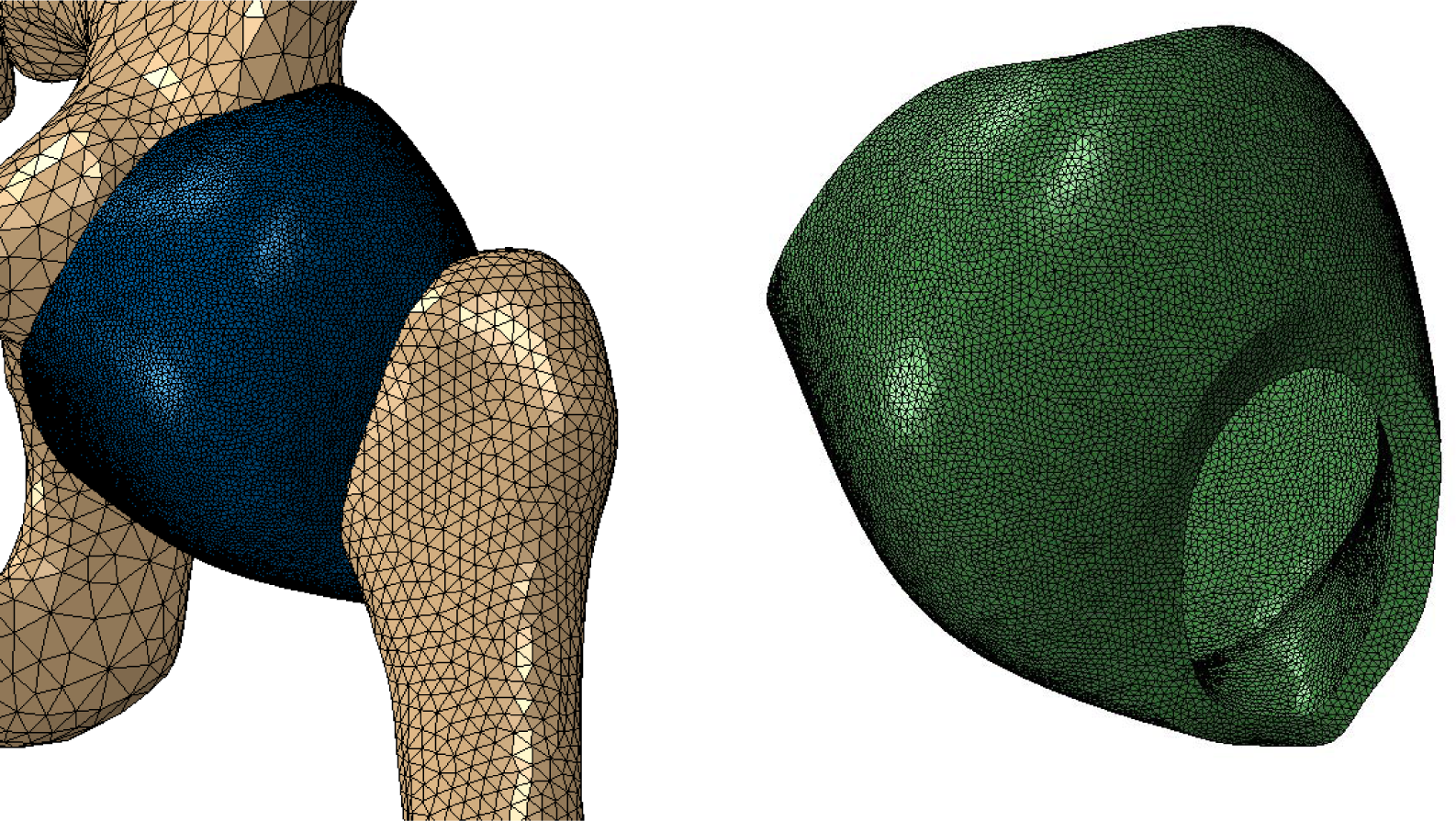
Hip capsule geometry and anatomical attachment sites based on Stewart et al [16].

A mesh convergence study was performed on the components of the model to ensure proper mesh density and quality. Meshlab [18] was used to apply surface mesh to each component which was converted into hybrid tetrahedral elements in Abaqus 2019 (Simulia, Providence, Rhode Island, USA). Mesh size was reduced to approximately half the total number of elements in varying iterations. The parts were subjected to the same loading and boundary conditions and the maximum von Mises stress was recorded. The mesh density was chosen by selecting the element size for each category that fell within 5% of the most densely meshed trial, as shown in Supplemental Table A. The meshed components were assembled in Abaqus 2019.

### Model Validation

Both linear elastic and hyperelastic material properties were used for the ligament and other soft tissue structures, while only linear elastic material properties were used to model the implant components and bony features as outlined in Supplemental Table 4.1 and 4.2 [15, 19]. In the knee joint, the anterior cruciate ligament (ACL), posterior cruciate ligament (PCL), medial collateral ligament (MCL), and lateral collateral ligament (LCL) were modeled with hyperelastic material models [20]. Peters et al [21] reported experimental stress-strain data of the knee ligaments for a range of ages in males and females. To age-match the material properties closely to the current model, the experimental data from a healthy 43-year-old female was utilized. A hyperelastic, neo-Hookean material model was applied to the ACL, PCL, MCL, and LCL [22]. The other soft tissue structures of the knee complex were modeled using truss elements with linear elastic material properties based on previous studies [23].

To validate the hip joint under traction forces, hip joint dislocation was compared against cadaveric hip load-displacement data [24]. The pelvis was fixed, and an axial load was applied distally on the femur, at the intercondylar fossa. Frictionless surface to surface interactions with exponential pressure-overclosure behavior were used to model the hip joint based on a previous FE study [25]. The hyperelastic, Yeoh material model was used to model the hip capsule as suggested by Stewart at el [16]. Yeoh material coefficients (Table 1) were experimentally determined to fit the curve of the reported cadaveric load-displacement data (Figure 4) [16, 24].

**Table 1.** Hyperelastic Yeoh material model coefficients for the hip capsule.

| **Yeoh**<br>**Coefficients** | C10 | C20 | C30 | D1 | D2 | D3 |
| --- | --- | --- | --- | --- | --- | --- |
|  | 0.65 | 0.75 | 6.85 | 0.000887 | 0.000487 | 0.000487 |

The stiffness of the knee ligament complex was validated against clinical data reported for ligament balancing using the Depuy Synthes Knee Balancer (Depuy Synthes, Warsaw, IN, USA) in PCR TKAs. For the validation study, the distal portion of tibia and fibula was encastered to restrict translation and rotation. Using SolidWorks, metal paddles resembling the Knee Balancer device were created and placed on the femoral condyles as directed by device protocol [26]. An axial force was applied on each paddle similar to the device mechanism described by Völlner et al [26]. Force displacement curves were constructed and compared against previously collected clinical data that used the same device [26, 27].

### Simulations

Three different models were assembled according to different TKA techniques performed: Bi-Cruciate Retaining (BCR) model, Posterior-Cruciate Retaining (PCR) model, and the Posterior Stabilized (PS) model (Figure 3A-C). The BCR model is noted by retaining all native ligaments of the knee joint (ACL, PCL, MCL, and LCL), whereas the PCR model was subject to ACL removal and the PS model required ACL and PCL removal [28]. The TKA instrumentation was modelled with a rigid “tie” interaction at the bone interface to limit motion between implant and bone. Frictionless, hard contact surface to surface interactions were applied at the patellofemoral and tibiofemoral joints. The femur was angled at 10LJ relative to the pelvis under the guidance of an orthopedic surgeon to fall within the range of previously reported positioning of the lower extremity [4, 29]. The pelvis was encastered to prevent its motion under the traction forces as motion of the patient’s trunk is restrained, intraoperatively [4]. To simulate the loading condition of hip distraction, an axial force was coupled to the distal fibula and tibia [29, 30]. Clinical studies have indicated a wide range of required traction forces (100N-600N) and a recommended minimum of 10 mm of hip joint distraction [8, 31–35]; thus, the present model was subjected to traction forces ranging between 100N to 500N to analyze joint behavior.

**Figure 3.**
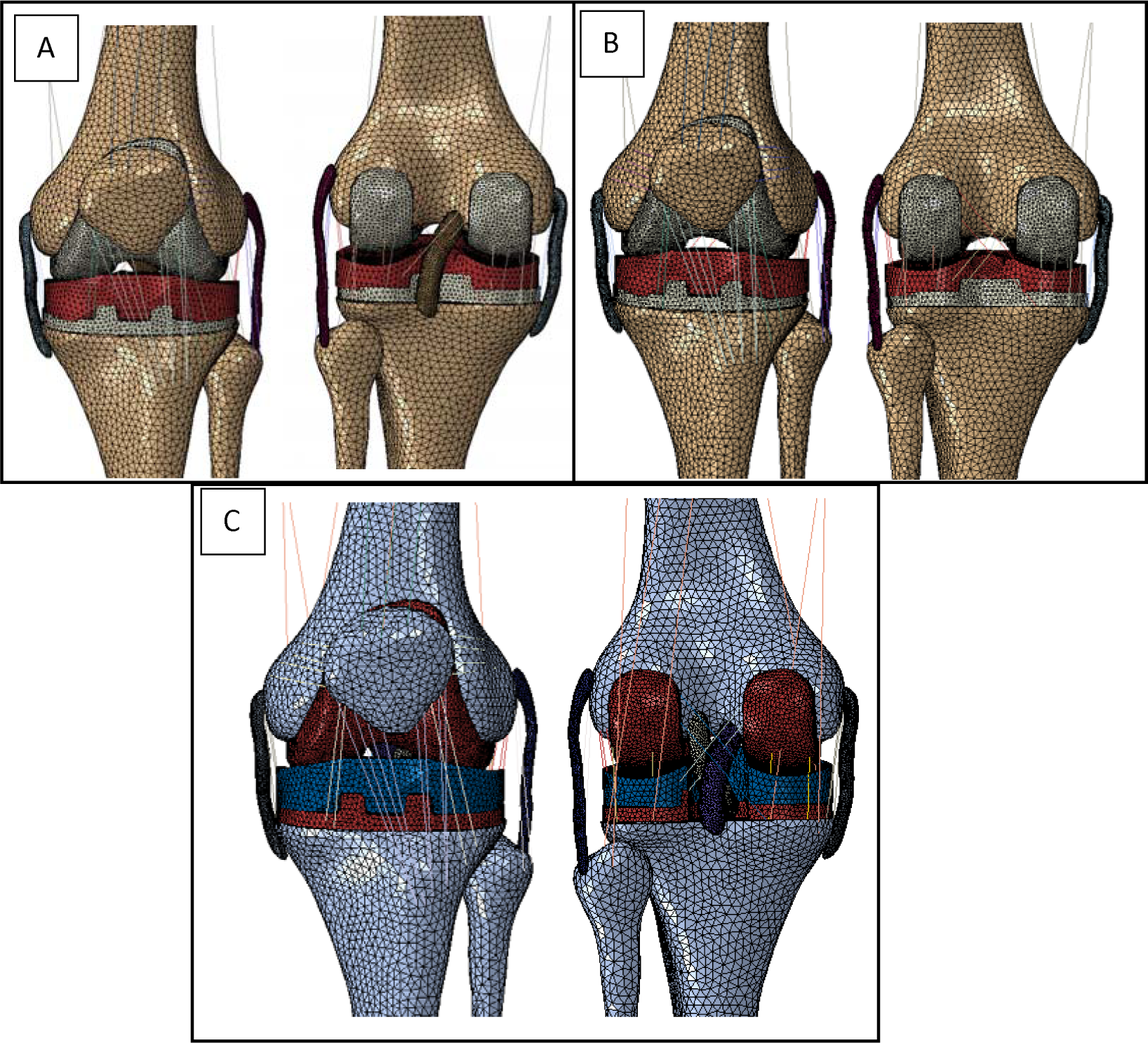
FE models of instrumented TKA. A) PCR TKA with simulated ACL release. B) PS TKA with simulated ACL and PCL releases. C) BCR TKA with intact ligaments.

### Data Analysis

Joint spacing and ligament strain were analyzed to assess the effects of traction forces. Relative strain (i.e. strain relative to pre-traction length) was calculated following methodology previously reported [36]. To measure the compartmental knee joint spacing, two points approximately vertical to each other were selected in the center region of the tray component and femoral component of the TKA system for the medial and lateral compartments. The difference before and after traction was measured to calculate the knee joint spacing. For hip joint distraction, a point selected on the femoral head near the centroid was used to calculate the displacement of the femur respective to the hip joint as suggested by a previous study [37].

### Results Validation

The FEA simulation of hip dislocation fell within an agreeable range of previously measured cadaveric data (Figure 4). Moreover, the joint stiffness for the PCR model fell within the ranges of clinical data shown in Table 2. The trend data for ligament balancing during a PCR TKA in Figure 4 fell within range of *in-vivo* data.

**Figure 4.**
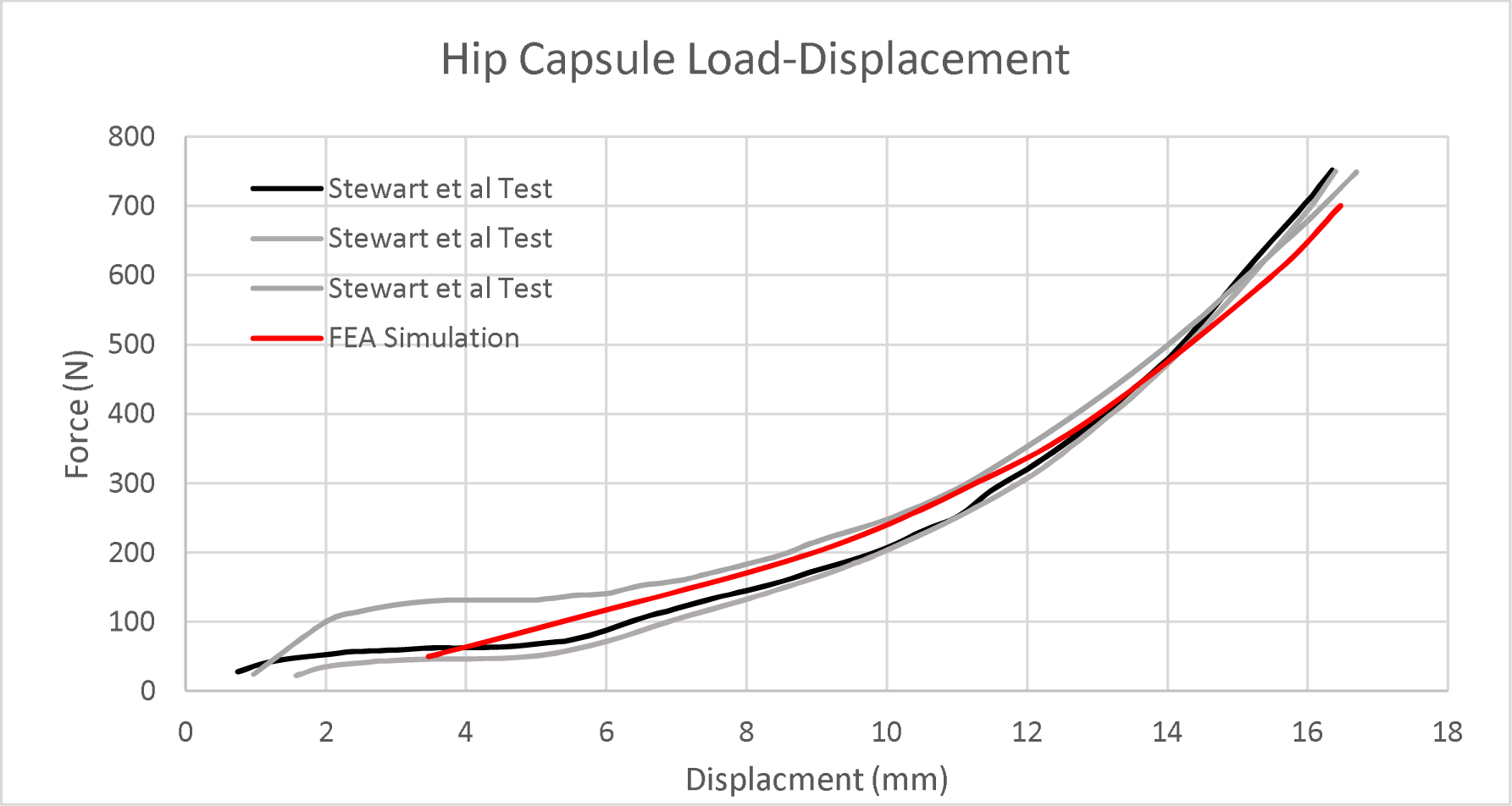
The load-displacement curve for the hip joint under various distraction forces. The gray lines represent experimental cadaveric data measuring hip joint dislocation [24]. The red line represents the FE model data with experimental curve fitted Yeoh hyperelastic material coefficients for the hip capsule.

**Figure 5.**
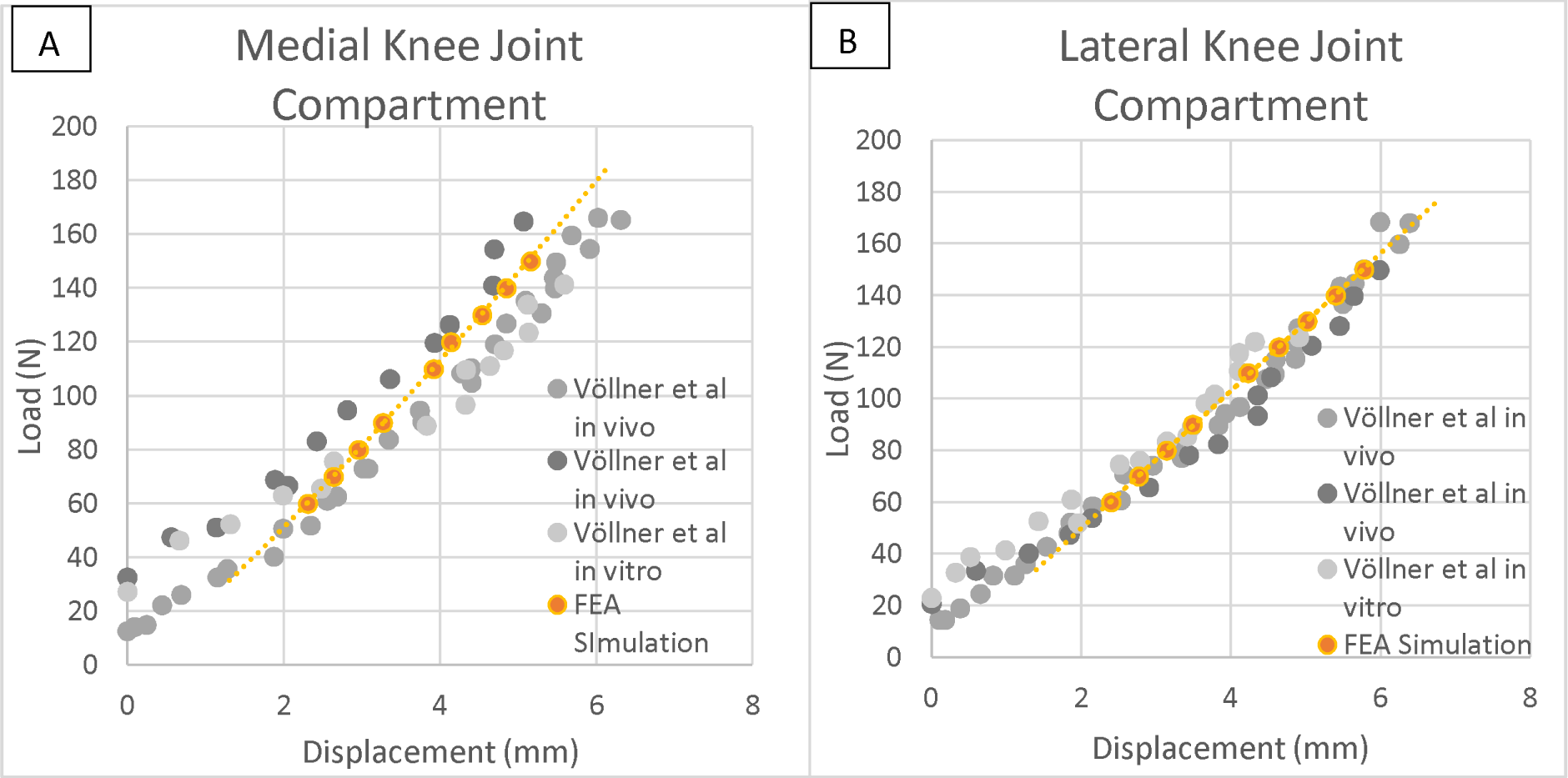
Knee ligament balancing for the A) medial and B) lateral compartment for PCR TKAs. The gray points represent cadaveric and in-vivo reported load-displacement values [26, 27]. The orange points represent the load-displacement data for the PCR model under knee ligament balancing.

**Table 2.** Knee compartmental stiffness for PCR TKAs during knee ligament balancing. The clinical measurements were taken with the DePuy Knee Balancer System (DePuy Synthes, Johnson and Johnson, Raynham, MA). The compartmental stiffness in the model was averaged across a range of simulated forces used in knee ligament balancing.

| **Compartmental Stiffness** |  |  |
| --- | --- | --- |
|  | lateral compartment<br>(N/mm) | Medial compartment<br>(N/mm) |
| **Cadaveric**<br>**[27]** | 31.6 | 26.6 |
| ***In vivo* [26,**<br>**27]** | 25.3 | 25.7 |
|  | 19.9± 0.9 | 28.4 ± 1.2 |
| **FE Model** | 25.7 | 27.8 |

### Hip Joint Distraction

Figure 6 displays the hip joint distraction for each of the implant scenarios. The implant scenario did not alter the force needed to distract the hip joint. Based on the clinical suggestion of 10 mm of hip joint spacing, adequate joint visualization for the present model was achieved approximately between 250 N to 350 N [37].

**Figure 6.**
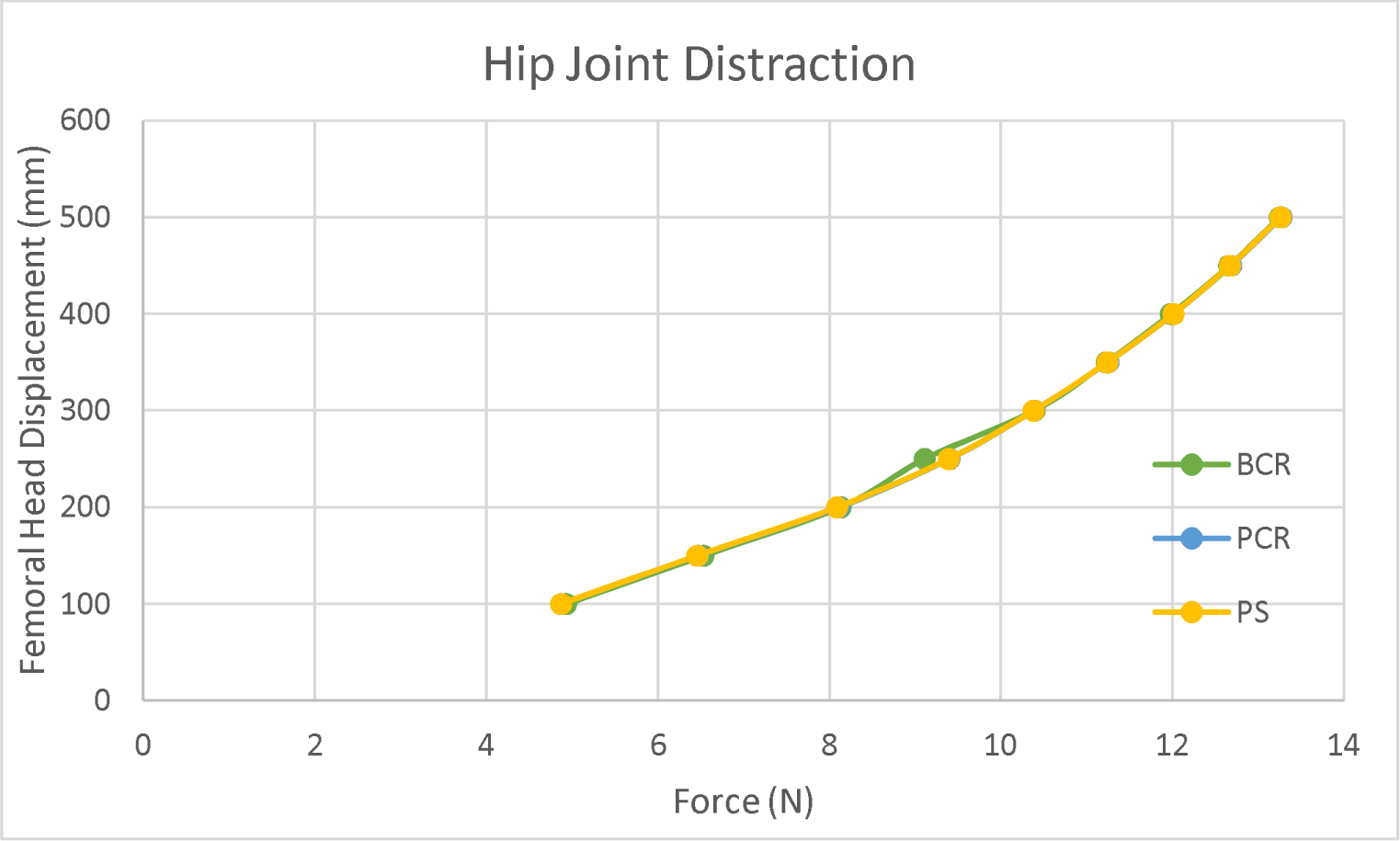
Hip joint distraction under axial traction force for the varying TKA implant scenarios.

### Knee Complex

The lateral and medial compartmental stiffness of the knee joint was analyzed under hip distraction for the three different TKA scenarios. The BCR model displayed the greatest average knee complex stiffness of 39.65 N/mm for the medial compartment and 37.48 N/mm for the lateral compartment compared to the PCR and PS implants (Figure 7). With release of the ACL for the PCR implant, there was a 31% and 34% decrease in the medial and lateral compartment stiffness, respectively, compared to the intact BCR model. Release of the PCL and ACL for the PS implant resulted in a stiffness decrease of 42% medially and 43% laterally compared to the BCR, but only a 16% and 15% decrease in medial and lateral compartment stiffness compared to the PCR case.

**Figure 7.**
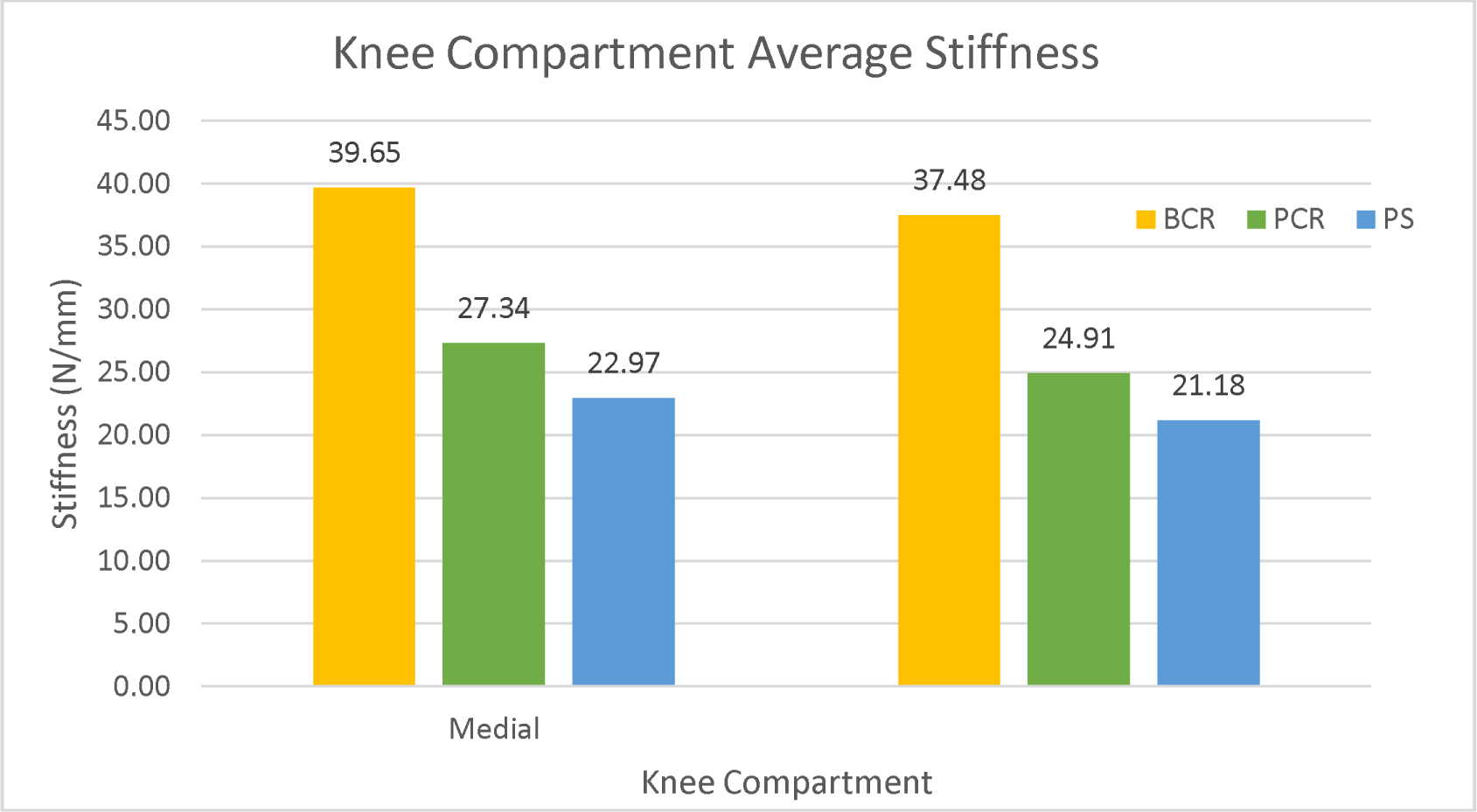
Averaged stiffness for the knee complex during the range of distraction forces.

The knee joint spacing during distraction was recorded. The PS implant resulted in the greatest knee joint spacing at each force, whereas the BCR model had the least amount of joint spacing for both compartments (Figure 8). On average, the PCR model’s medial compartment resulted in an average of 45% greater joint spacing compared to the BCR model, and the PS model resulted in a 72% increase in medial compartment spacing.

**Figure 8.**
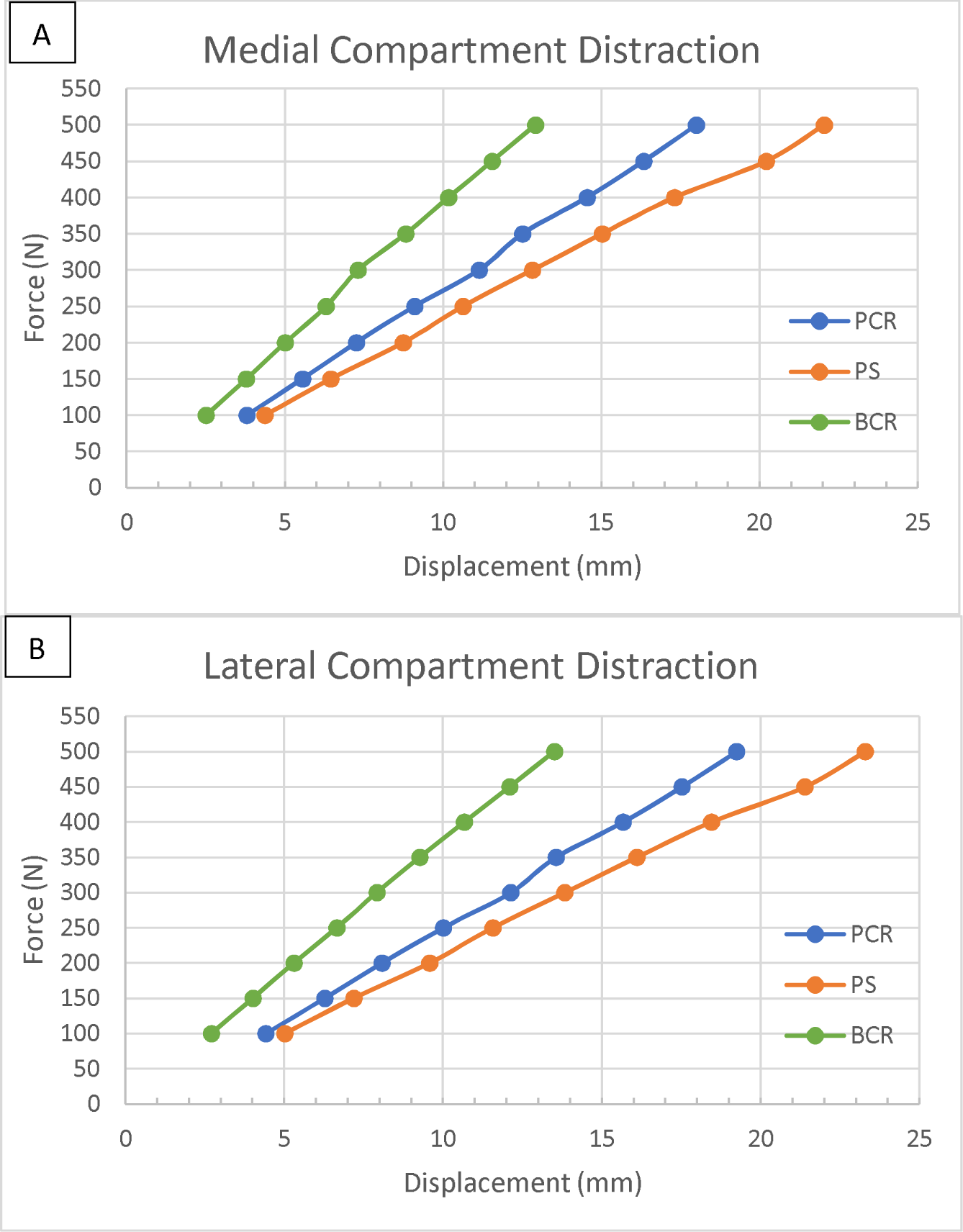
Knee joint distraction under the axially applied traction force for the A) medial and B) lateral compartment.

### Knee Ligament Strain

The relative strain of the four main ligaments were measured under traction forces. The MCL ligament displayed the highest levels of strain in the PS model and the lowest strains in the BCR model. Comparing the PCR and PS cases, the presence of the PCL played a larger role in reducing MCL strain compared to the LCL, especially at higher loading conditions (Figure 9). The ACL was noted to have the lowest strain out of the ligaments present in the BCR model. The presence of the ACL played a greater role in reducing MCL and LCL strain compared to the PCL. The percent increase in strain of the MCL and LCL for the PCR model compared to the BCR model was 54% and 67%, respectively. The percent increase in strain for the MCL and LCL for the PS model compared to the PCR model was 18% and 15%, respectively. However, the strain for the MCL and LCL for the PS model compared to the BCR model increased by 87% and 95%, respectively.

**Figure 9.**
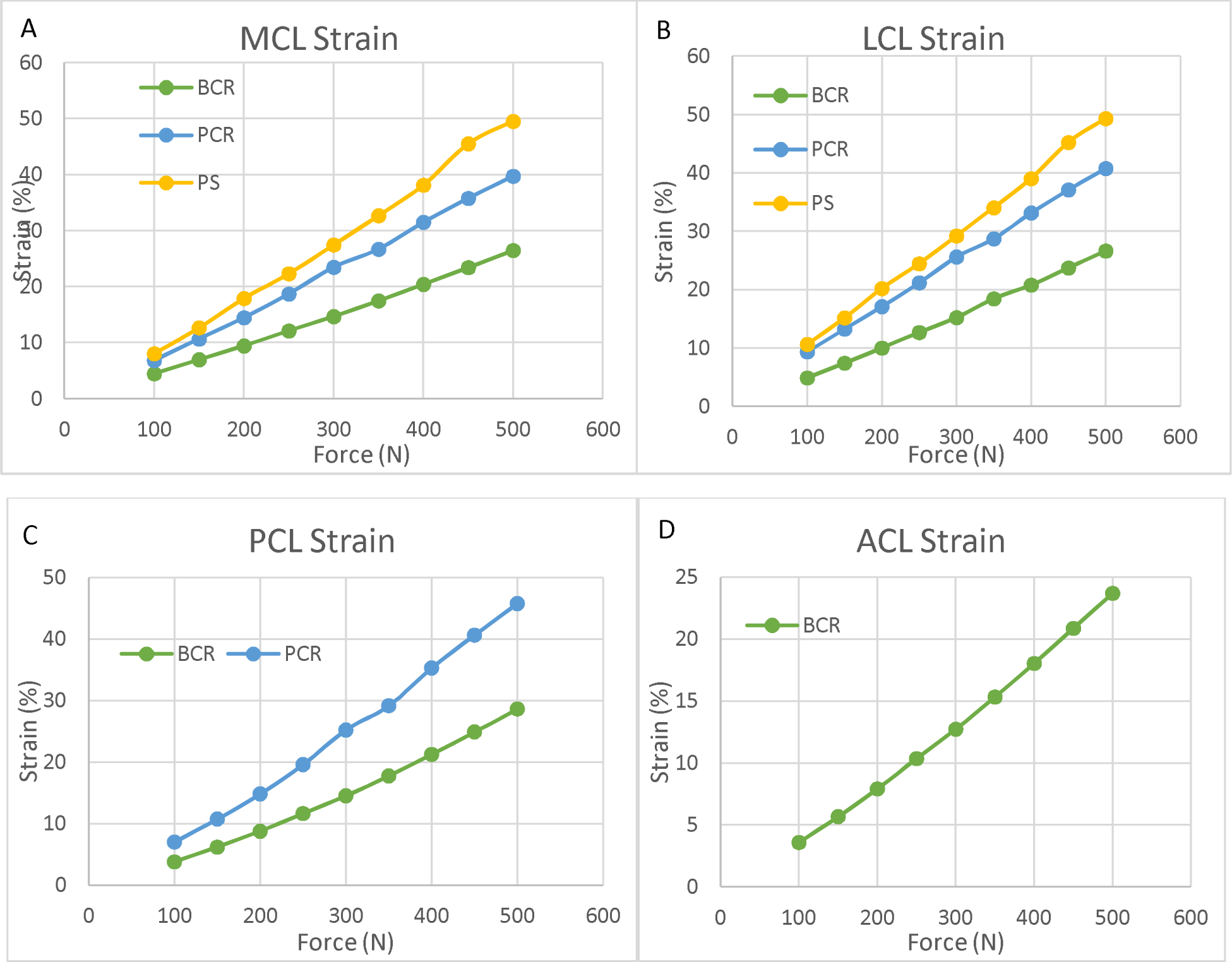
Measured strain in the A) MCL, B) LCL, C) PCL, and D) ACL under the traction forces.

## Discussion

Our findings indicate that knee joints post-surgery, particularly those with ligament release, may face a heightened risk of further ligamentous damage due to excessive distraction. The PCR knee and PS model were subject to excess knee joint distraction and ligament strain before adequate hip joint distraction was achieved. The bi-cruciate retaining TKA was indicated to be the safest under the distraction conditions, however strain levels were indicative of potential soft tissue injury, which supports the common clinical reports of pain and instability in the knee joint, post-operatively. This suggests a need for the development of safer distraction techniques to minimize knee joint loading during hip distraction in vulnerable knee joints.

The reported complication rates for hip arthroscopies vary from 1.4% to 7%, encompassing issues such as transient neurapraxia and iatrogenic soft tissue injury [38–41] largely attributed to the substantial forces required for hip joint distraction [7, 40]. Despite awareness of these risks, research into the biomechanical impact of distraction forces on the knee joint during surgery remains scarce [6]. Beutel et al [42] investigated hip arthroscopy complications in a small cohort of 9 patients with previous lower-extremity arthroplasty, finding no adverse effects. However, with only five of these patients having undergone knee arthroplasty (either total or uni-compartmental), the findings offer limited generalizability. This study, therefore, seeks to fill a critical gap by examining the biomechanical responses of knee joints with prostheses when subjected to traction forces, aiming to enhance the safety and efficacy of hip arthroscopy techniques.

The results of the validation simulations agreed with previous literature findings for knee ligament complex stiffness in patients with posterior cruciate retaining TKRs. The hip joint dislocation under axial traction forces were similar to cadaveric load displacement curves (Figure 4). Recognizing the variability in patient-specific traction forces [31], the current study utilized a wide range of forces to account for these differences. Recent research has identified a positive correlation between traction forces and factors such as body weight and the presence of hip osteoarthritis [31]. However, Yin at el suggests lower extremity muscle volume is a greater indicator of higher traction forces than BMI [43]. Thus, these patient factors must be considered when determining risk for patients with prior TKAs.

The stiffness of the hip joint (Figure 4) was shown to be greater than that of the knee complex subjected to PCR and PS TKAs (Figure 7). Previous literature supports these findings with the hip capsule stiffness ranging between 12.71 N/mm to 60.20 N/mm based on section [24]. However, there is limited knowledge of the intact stiffness of the knee complex under tensile forces [44]. Our results from the BCR model with intact knee ligaments report a stiffness of 39.65 N/mm and 37.78 N/mm for the medial and lateral compartments, respectively. Based on the previous literature findings for hip joint stiffness and our model predictions, the lower stiffness of the knee joint may predispose it to negative effects from hip distraction forces. Moreover, we observed a large reduction in knee complex stiffness following ligament releases for PCR and PS TKAs (Figure 7). Ligament balancing, aimed at correcting varus and valgus deformities in TKR patients, often involves sequential ligament releases in PCR TKAs. Yet, literature advises against complete MCL releases due to its critical role in passive knee stabilization [44] with stiffness potentially decreasing by up to 40% post-release [44]. Based on the trends outlined in the present study, patients undergoing additional ligament releases may be at an increased risk for injury during hip distraction. The contribution of the PCL to knee stability under tensile forces remains a topic of debate, with no consensus on its biomechanical role when the knee is fully extended [44, 45]. Figure 8 suggests a relationship between the PCL’s contribution to stability and the applied traction force. Our results indicate that the PCL’s role in passive stabilization is comparatively minor, with only a 15% and 16% stiffness difference observed between the PCR and PS models, respectively. This underscores the predominant stabilizing functions of the ACL and MCL.

Knee complex stiffness was observed to vary with changes in intraoperative knee joint spacing and ligament strain, showing an average increase in joint spacing of 45% for PCR and 72% for PS models relative to the intact BCR model (Figure 8). While injury can occur within a wide range of strain values depending on patient specific factors, there is limited data quantifying ligament strains in intact knee kinematics [46, 47]. In the present study, within the traction force ranges of 250N to 350N needed for adequate joint visualization, the strain for the MCL and LCL was greater than 20% for the PCR and PS simulations which may suggest potential for complications [46]. The BCR model resulted in strain levels that remained below 20%. Moreover, when knee joint spacing exceeds 10 mm of distraction there may be risk for ligament failures and iatrogenic dislocation due to high strain values noted in Figure 9. BCR implants have been debated due to poor surgical outcomes with unclear indications for use, making it a less widely used implant compared to the PCR [48]. In one study of a small cohort, only 16% of patients received a BCR with the main indication of use being an intact ACL [49]. The present study suggests BCR implants may reduce the chance of traction related complications because of retaining the native ligaments. The strain values noted in the BCR model were within reasonable range compared to strain values reported during native knee motion, suggesting a minimal risk of injury [50, 51]. However, when surpassing the recommended 10 mm of distraction, the strain levels in the ACL suggest minor injury may occur. Our FE data findings, which show large changes in knee complex stiffness and joint spacing, align numerically with clinical observations reported in the literature, where approximately 50% of patients experience sensations of instability and pain in the knee following procedures involving hip distraction [6]. This numerical agreement suggests a potential biomechanical underpinning for the clinical symptoms observed post-procedure, warranting further investigation to explore the direct impact of biomechanical alterations on patient outcomes.

The high risk of traction related problems has been recognized, and in response, various surgical procedures have emerged to reduce traction force and time [52, 53]. Lateral or supine patient positioning are used during hip arthroscopy, but the superiority of one over the other is debated [54]. Despite the approach used, abduction and flexion of the hip are used to relax the hip capsule to allow for easier distraction [10, 55, 56]. Additionally, various surgical interventions have been proposed to alter the resistive forces of the hip capsule. Distension combined with traction increases joint distraction upwards of 2.25 times compared to traction alone [57, 58]. Furthermore, joint venting prior to the application of traction significantly reduces the amount of traction needed, however, the required force still fell within the range of 100 N to 450N [34, 59]. Based on the trend outlined in the present study, the force required remains in a harmful range for patients with PCR and PS TKRs. Moreover, capsulotomies have been a debated procedure to reduce traction forces [60, 61]. The size of the capsulotomy is known to reduce the traction force; however, capsule repair is essential to return the capsule to its native functionality [62, 63]. The combination of joint venting and capsulotomies further reduces the traction force safely, but the lower extremity joints remain exposed to excessive traction [8, 34]. Further research is needed to study and combat hip distraction forces for vulnerable patient populations.

### Limitations

While our findings underscore the pivotal role of the ACL in knee stabilization during distraction forces, it’s important to note that the material properties used in our analysis were derived from a healthy, non-arthritic individual. Given the known susceptibility of intraarticular ligaments to degradation from osteoarthritis (OA) and aging [21, 64] our study’s application of semi-patient-specific material properties may not fully encapsulate the biomechanical realities of younger patients or those with advanced OA. Despite this limitation, our results indicate a discernible pattern: individuals with a history of TKA, potentially exhibiting compromised ACL and PCL properties due to OA, could be at an elevated risk of injury during surgery. This insight, while derived from a constrained model, highlights the necessity for further research into the biomechanical implications of ligament degradation on surgical outcomes.

Our approach to modeling muscle forces as uniaxial truss-connector elements with linear, elastic material properties, while simplified [19], closely mirrors the clinical scenario during hip distraction procedures where patients are under general anesthesia. In this state, muscles indeed function as passive stabilizers, akin to the passive mechanical behavior assumed in our simulations. This aspect of our model is particularly relevant given that, under anesthesia, the dynamic and active properties of muscle forces are substantially reduced, aligning with the conditions we have simulated. A previous study highlighted increased hip traction forces in awake patients [32], a situation divergent from standard practice where anesthesia is employed, thus reinforcing the clinical applicability of our modeling assumptions. Therefore, rather than detracting from the study’s validity, this simplification may, in fact, enhance the relevance of our findings to the typical clinical environment, where the procedure is performed under conditions that render muscle forces passive. This alignment with clinical practice supports the legitimacy of our study and highlights the potential applicability of our results in understanding the biomechanical implications of hip distraction procedures.

Lastly, the PS model was simplified by using the PCR model with a PCL release. While simplified in the present study, a PS implant commonly has a uniquely designed cam-post that limits anterior-posterior translation. However, this would not alter the mechanism under traction forces [65].

### Conclusions

While traction forces are a significant concern for patient outcomes in hip distraction procedures, the detailed biomechanical impact on the lower extremity, particularly following TKA, has not been fully explored. Our biomechanical models suggest that individuals who have undergone hip arthroscopy post-TKA with ligament release might be at a heightened risk for subsequent soft tissue damage, highlighting the importance of carefully evaluating a patient’s TKA history when assessing the risks associated with hip distraction. Because of the inherent instability present in TKAs, post-operatively, surgeons should exercise increased caution and incorporate techniques to lower traction forces for patients with prior knee ligament releases. The insights from our study point towards the necessity of further research into the biomechanical consequences of hip distraction in patients with previous TKAs, aiming to refine safety protocols and enhance patient care.

## Supplemental Data

### Mesh Convergence

A component from each of the bones, implant, and ligaments geometries, were utilized in the mesh convergence study to appropriately assign mesh density. The maximum Von Mises stress was recorded for varying mesh sizes. For the bone components (Supplemental Table 1), a mesh size of 3 was selected. A mesh size of 1 was selected for the 3D soft tissue components (ligaments and hip capsule) (Supplemental Table 2). Moreover, a mesh size of 2 was selected for each of the TKA implant components (Supplemental Table 3).

**Supplemental Table 1.** Mesh convergence study of the femur under tensile load of 50N.

| # of Elements | Mesh Size | Max Von Mises (MPa) |
| --- | --- | --- |
| 297541 | 2 | 1.851 |
| 225953 | 2.25 | 1.827 |
| 173626 | 2.5 | 1.824 |
| 125185 | 2.75 | 1.8 |
| 100724 | 3 | 1.782 |

**Supplemental Table 2.** Mesh convergence study of the LCL ligament under tensile load of 50N.

| # of Elements | Mesh Size | Max Von Mises (MPa) |
| --- | --- | --- |
| 17285 | 0.75 | 3.703 |
| 7495 | 1 | 3.695 |
| 4034 | 1.25 | 3.683 |

**Supplemental Table 3.** Mesh convergence study of the tibial component under tensile load of 50N.

| # of Elements | Mesh Size | Max Von Mises (MPa) |
| --- | --- | --- |
| 126727 | 1 | 2.888 |
| 77747 | 1.25 | 3.013 |
| 46128 | 1.5 | 3.072 |
| 29369 | 1.75 | 3.099 |
| 20793 | 2 | 3.034 |

### Material Properties

**Supplemental Table 4.1.** Implant material properties [15].

| **Part** | **Material** | **Young's Modulus (MPa)** | **Poisson's Ratio** |
| --- | --- | --- | --- |
| Femoral Component | Titanium | 120,000 | 0.3 |
| Tibial Tray | Polyethylene | 690 | 0.46 |
| Tibial Component | Titanium | 120,000 | 0.3 |
| Patellar Dome | Polyethylene | 690 | 0.46 |

**Supplemental Table 4.2.** Hard tissue material properties [15].

| **Material** | **Young's Modulus (MPa)** | **Poisson's Ratio** |
| --- | --- | --- |
| Cortical Bone | 15,000 | 0.3 |
| Cancellous Bone | 400 | 0.3 |

